# Connecting gene regulatory relationships to neurobiological mechanisms of brain disorders

**DOI:** 10.1101/681353

**Authors:** Nancy Y. A Sey, Harper Fauni, Won Ma, Hyejung Won

**Affiliations:** UNC Neuroscience Center, University of North Carolina, Chapel Hill, NC 27599, USA; Department of Genetics, University of North Carolina, Chapel Hill, NC 27599, USA

## Abstract

Despite being clinically distinguishable, many neuropsychiatric disorders display a remarked level of genetic correlation and overlapping symptoms. Deciphering neurobiological mechanisms underlying potential shared genetic etiology is challenging because (1) most common risk variants reside in the non-coding region of the genome, and (2) a genome-wide framework is required to compare genome-wide association studies (GWAS) having different power. To address these challenges, we developed a platform, *Hi-C coupled MAGMA* (*H-MAGMA*), that converts SNP-level summary statistics into gene-level association statistics by assigning non-coding SNPs to their cognate genes based on chromatin interactions. We applied H-MAGMA to five psychiatric disorders and four neurodegenerative disorders to interrogate biological pathways, developmental windows, and cell types implicated for each disorder. We found that neuropsychiatric disorder-associated genes coalesce at the level of developmental windows (mid-gestation) and cell-type specificity (excitatory neurons). On the contrary, neurodegenerative disorder-associated genes show more diverse cell type specific, and increasing expression over time, consistent with the age-associated elevated risk of developing neurodegenerative disorders. Genes associated with Alzheimer’s disease were not only highly expressed in microglia, but also subject to microglia and oligodendrocyte-specific dysregulation, highlighting the importance of understanding the cellular context in which risk variants exert their effects. We also obtained a set of pleiotropic genes that are shared across multiple psychiatric disorders and may form the basis for common neurobiological susceptibility. Pleiotropic genes are associated with neural activity and gene regulation, with selective expression in corticothalamic projection neurons. These results show how H-MAGMA adds to existing frameworks to help identify the neurobiological basis of shared and distinct genetic architecture of brain disorders.

## Introduction

Psychiatric disorders display shared symptoms and comorbidities, which makes psychiatric nosology difficult. This is in stark contrast with neurodegenerative disorders, which have greater diagnostic specificity. Understanding the genetic relationship between different neuropsychiatric illnesses can complement current diagnosis frameworks based on clinical presentations by informing the genetic basis of psychiatric disorders. Investigators have developed statistical frameworks, such as polygenic risk scores and cross-trait LD score regression, to estimate genetic correlations for various traits based on genome-wide SNPs^1^. Indeed, recent cross-trait genetic correlation analyses demonstrate widespread shared genetic etiology among psychiatric disorders, but not in neurological disorders^2^. However, it is still challenging to decipher the shared or distinct neurobiological mechanisms, largely because the majority of common risk variants reside in non-coding regions of the genome^3^. A common practice is to assign risk variants to the nearest gene or to use linkage disequilibrium information, but neither of these is accurate^4–6^.

Since genes do not act alone, but act in pathways, and individual common variants even when collapsed on a single gene have limited statistical power to show association with a disorder, several tools have been developed to collapse association analyses onto biological relevant pathways^7–11^. One of the most commonly used tools, Multi-marker Analysis of GenoMic Annotation (MAGMA) was developed to aggregate SNP associations into gene-level associations while correcting for many confounding factors such as gene length, minor allele frequency, and gene density^7^. MAGMA is a powerful tool that has been used broadly, and can (1) pinpoint genes that are strongly associated with a given trait, and (2) reduce the multiple testing correction burden that allows the detection of signals in non-genome-wide significant (GWS) loci. However, MAGMA and other pathway tools do not take long-range regulatory interactions into consideration, largely disregarding the majority of intergenic variants and assigning intronic variants to the genes in which they reside.

It is becoming increasingly recognized that long range (>10kb) regulatory interactions are related to 3D chromatin structure, whereby distal enhancers are brought into contact with the gene promoter^6,12^. Hi-C identifies genome-wide chromatin configuration, which provides a framework for assigning non-coding variants to genes. We therefore modified MAGMA approach to create *Hi-C coupled MAGMA* or *H-MAGMA*, that leverages Hi-C datasets to assign non-coding SNPs to their cognate genes. We applied this framework to generate gene-level summary statistics for five neuropsychiatric disorders (Attention deficit hyperactivity disorders, ADHD; Autism spectrum disorders, ASD; Schizophrenia, SCZ; Bipolar disorder, BD; Major depressive disorders, MDD) and four neurodegenerative disorders (Amyotrophic lateral sclerosis, ALS, Multiple sclerosis, MS; Alzheimer’s disease, AD, and Parkinson’s disease, PD, Figure 1). H-MAGMA identified more significantly associated genes than conventional MAGMA by incorporating non-coding SNPs during the conversion of SNP-level association statistics into gene-level association statistics. Gene-level association statistics from H-MAGMA closely resemble genetic relationships among brain disorders, which enable subsequent analyses to identify biological pathways, developmental windows, and cell types critical for each brain disorder.

**Figure 1.**
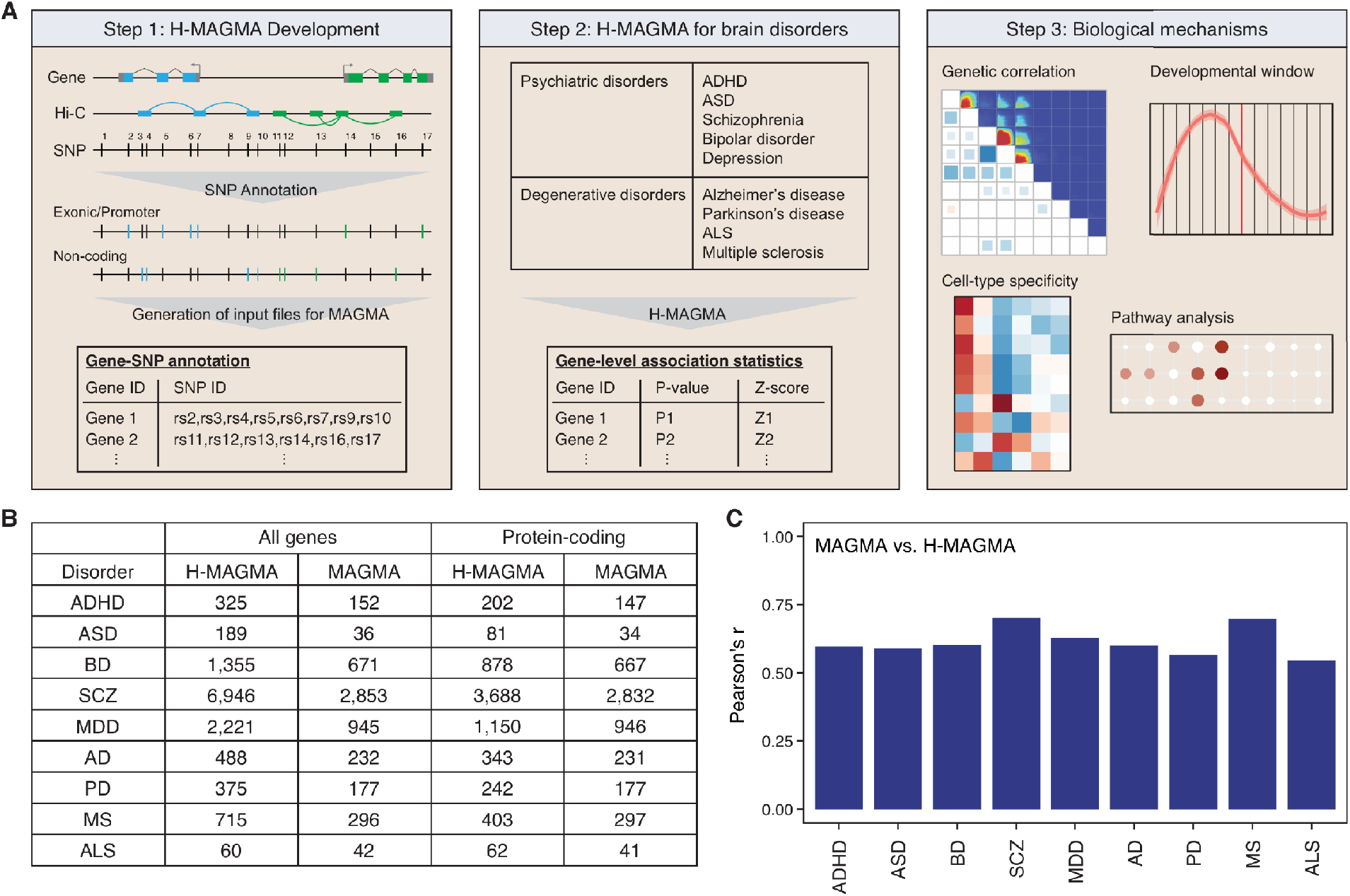
Schematics of H-MAGMA approach. **A.** H-MAGMA leverages chromatin interactions (Hi-C) to assign intergenic and intronic SNPs to putative genes. We have applied this framework for five psychiatric disorders and four neurodegenerative disorders by leveraging Hi-C datasets derived from the fetal and adult brain. H-MAGMA gives gene-level association statistics, which can be used to identify biological mechanisms underlying brain disorders. **B.** By incorporating intragenic SNPs, H-MAGMA identifies more significantly associated genes (FDR<0.05) than conventional MAGMA (MAGMA). **C.** The gene-based association statistics of conventional MAGMA and H-MAGMA are moderately correlated.

## Result

### Hi-C coupled MAGMA

Since our primary goal is to identify neurobiological mechanisms underlying brain disorders, we leveraged two Hi-C datasets obtained from human brain tissue, one from the fetal brain^4^ and the other from the adult brain^13^. Using Hi-C datasets, we generated gene-SNP pairs that reflect long-range regulatory relationships, which serves as an input file for MAGMA (Figure 1, **Supplementary Data File, Methods**). Exonic and promoter SNPs were directly assigned to their target genes based on their genomic location, while intronic and intergenic SNPs were assigned to their cognate genes based on chromatin interactions (Figure 1A). When assigning SNPs to genes, we used chromatin interactions from promoters and exons, as our previous study comparing Hi-C data with expression quantitative trait loci (eQTLs) have demonstrated the gene regulatory potential of exon-level interactions^13^.

This framework advances conventional MAGMA as it increases power of detecting gene-level associations by leveraging previously discarded intergenic SNP association statistics, and it correctly assigns non-coding SNPs to the putative target genes based on functional genomics evidence. It also expands functional genomics-guided gene pruning for GWS loci^4,13,14^, by utilizing genome-wide association statistics rather than restricting the test within genome-wide significant loci. By expanding the search space to sub-threshold loci, it enables (1) pathway analyses for relatively low-powered genome-wide association studies (GWAS), and (2) comparing GWAS with different sample sizes and different numbers of GWS loci.

We applied this framework to nine brain GWAS, including five neuropsychiatric disorders and four neurodegenerative disorders (Figure 1A, **Supplementary Table 1**). As expected, the number of genes significantly associated with brain disorders (FDR<0.05) was increased compared with conventional MAGMA, supporting the increase in power by incorporating intergenic signals (Figure 1B, **Supplementary Table 2**). Gene-level association statistics were modestly correlated between conventional MAGMA and H-MAGMA (correlation coefficient=0.6-0.7), indicating that leveraging functional genomics in annotating non-coding variation can identify novel genes and pathways (Figure 1C).

### Psychiatric disorders exhibit neurodevelopmental origin, while degenerative disorders exhibit adult origin

Since 3D chromatin loops are highly tissue-specific^4^, it is important to decide which Hi-C datasets are appropriate to use to identify target genes for each disorder. To address this, we first measured the heritability enrichment of each disorder using tissue-specific regulatory elements (**Methods**)^15,16^. Notably, psychiatric disorders showed strong enrichment in brain tissues, while neurodegenerative disorders lacked brain-specific enrichment (Figure 2A, **Supplementary Figure 1**). Among neurodegenerative disorders, PD was the only disorder that showed modest brain enrichment, while AD and MS showed strong enrichment in immune cells. This is in line with the growing evidence that MS and AD have strong immune basis^17–21^. Within brain tissues, psychiatric disorders showed stronger heritability enrichment in the fetal brain than in the adult brain, highlighting their neurodevelopmental origin. Fetal enrichment was more robust in neurodevelopmental disorders such as ADHD and ASD than in adult onset disorders including BD, SCZ, and MDD. On the contrary, neurodegenerative disorders showed stronger enrichment of heritability in adult brains.

**Figure 2.**
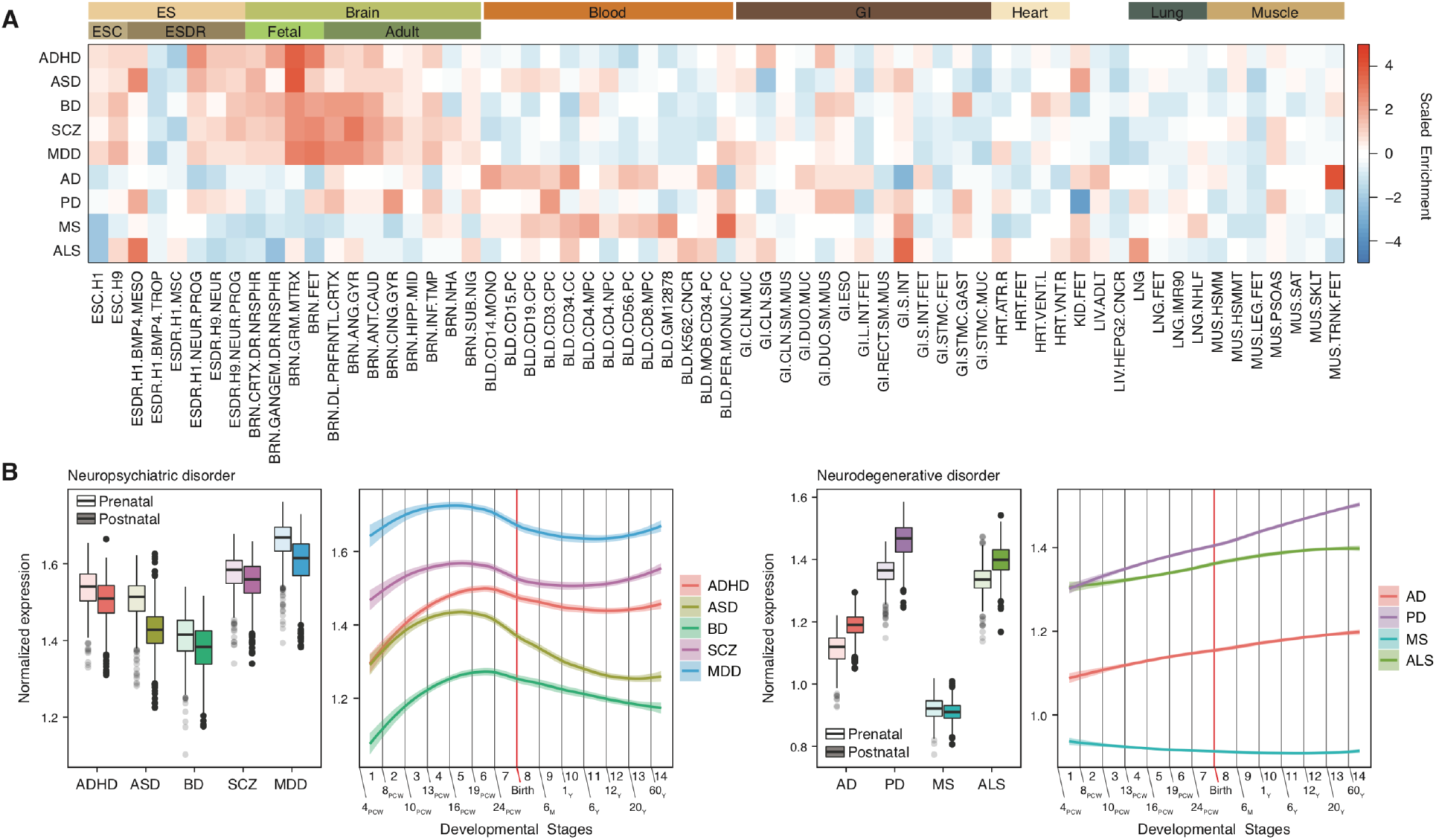
Spatiotemporal dynamics of brain disorder common risk genes. **A.** Heritability enrichment of brain disorders in active regulatory elements of multiple tissue/cell types suggests that psychiatric disorders show brain-selective enrichment, while neurodegenerative disorders lack brain enrichment. ESC, embryonic stem cells; ESDR, embryonic stem cell derived cells; GI, gastrointestinal. Abbreviations for each tissue/cell type are described in Methods. **B.** Developmental expression trajectories of brain disorder-associated genes. PCW, post-conception week; M, month; Y, year. (Left) N = 410 and 453 for prenatal and postnatal samples, respectively. Center, median; box=1st-3rd quartiles (Q); lower whisker, Q1 - 1.5 x interquartile range (IQR); upper whisker, Q3 + 1.5 x IQR. (Right) LOESS smooth curve with 95% confidence bands.

To confirm this result based entirely on regulatory enrichment, we also used an alternative gene-centric approach. Genes associated with each brain disorder were identified based on fetal and adult brain H-MAGMA (Figure 1A-B, **Methods**). We then compared expression values of disorder-associated genes between prenatal and postnatal stages (Figure 2B). There was a clear distinction between psychiatric and neurodegenerative disorders. Genes associated with psychiatric disorders were highly expressed during prenatal stages, while genes associated with neurodegenerative disorders were most highly expressed postnatally (Figure 2B, see **Supplementary Table 3** for statistics). The only exception was MS, as MS-associated genes displayed prenatal enrichment (FDR=2.94×10^−8^).

Next, we plotted the developmental expression trajectories for genes associated with each brain disorder (Figure 2B). ASD-, SCZ-, and MDD-associated genes showed remarkedly similar expression patterns with a peak at the developmental stage 5 (16-19 PCW)^22^. BD- and ADHD-associated genes were gradually increased during the prenatal stage with a peak at the developmental stage 6 (19-22 PCW). Developmental stages 5 and 6 both represent mid-gestation, the period during which upper layer neurons are generated and neuronal differentiation including axonogenesis and dendritic arborization takes place^22,23^. Therefore, this result collectively highlights mid-gestation as a critical window during neurodevelopment that may confer risk to multiple psychiatric disorders, consistent with recent results from cross-disorder GWAS^24,25^. On the contrary, neurodegenerative disorders showed a variety of distinct expression trajectories. AD-, PD-, and ALS-associated genes constantly and gradually increased during both prenatal and postnatal stages, suggesting that these genes may become more susceptible upon aging. MS-associated genes displayed enrichment during the first trimester. This is consistent with a strong neurodevelopmental predisposition for psychiatric disorders, in contrast with neurodegenerative disorders, which have a postnatal origin.

### Shared genetic architecture among brain disorders

We next assessed whether the gene-level association statistics obtained from H-MAGMA can be used to elucidate the shared genetic relationship among brain disorders. Since the number of genes significantly associated with a given disorder differs based on the sample size and power of GWAS, we used a rank-rank hypergeometric test of overlap (RRHO), which is a threshold-free algorithm for comparing two genomic datasets^26^. Genes were ranked based on Z-scores from the H-MAGMA output (Figure 1B), and ranked lists between two disorders were compared to identify the gene-level overlap between two disorders (**Methods**). We then compared this gene-level overlap with genetic correlations calculated by LD score regression (LDSC)^1^.

We were able to recapitulate the previously reported genetic architecture of brain disorders – that psychiatric disorders are genetically correlated while neurologic disorders are genetically distinct^2^. More importantly, this genetic architecture was also reflected in the gene-level overlaps. Psychiatric disorders exhibited strong overlaps in their ranked lists, whereas neurodegenerative disorders did not display significant overlaps (Figure 3A, **Supplementary Figure 2**). In particular, ADHD and ASD showed a strong overlap, and BD, SCZ, and MDD showed strong overlaps, suggesting shared biological pathways among neurodevelopmental disorders, as well as adult-onset psychiatric disorders. The correlation between RRHO and genetic correlation was 0.79 (Figure 3B, P-value=8.08×10^−9^), demonstrating that gene-level association statistics from H-MAGMA reflect shared genetic architecture, and hence can be further used to decipher the biological mechanisms underlying shared genetic architecture among brain disorders.

**Figure 3.**
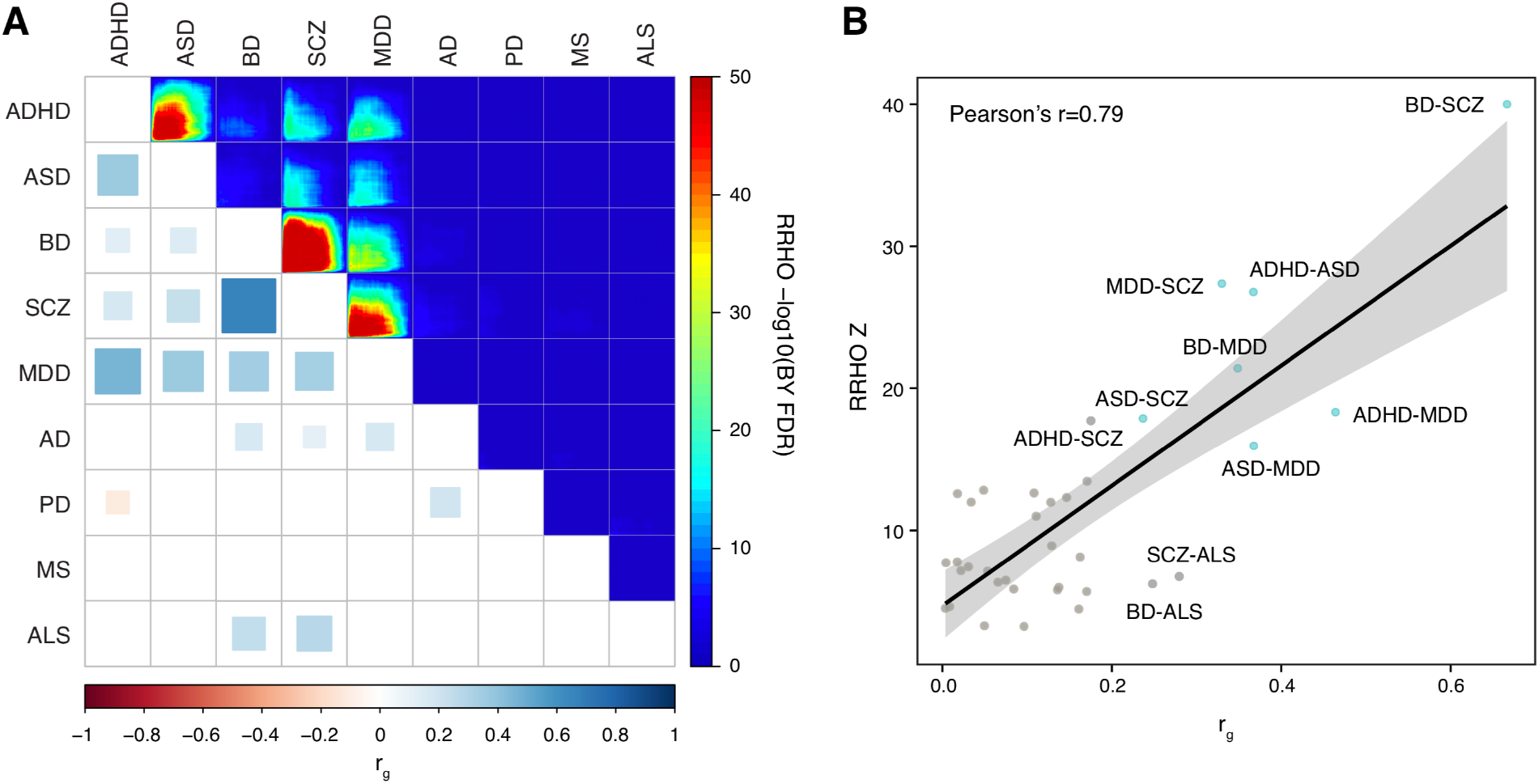
Genetic relationships among brain disorders. **A.** Psychiatric disorders show strong genetic relationships both at the level of genetic correlations (bottom left, r_g_) and gene-level overlaps (top right, RRHO). BY FDR, FDR using the Benjamini and Yekutieli procedure. **B.** Genetic correlations (r_g_) and gene-level overlaps (RRHO Z) are highly correlated, indicating that gene-level overlaps obtained by H-MAGMA recapitulate genetic correlations. Brain disorders that show strong genetic correlations (r_g_ > 0.2) and gene-level overlaps (RRHO Z > 15) are marked in blue. Linear regression line with 95% confidence bands.

### Pathways implicated for brain disorders

To identify biological pathways underlying psychiatric and neurodegenerative disorder risk, we conducted gene ontology analysis on gene-level association statistics from H-MAGMA. We again ranked genes based on Z-scores so that genes with higher Z-scores (more significantly associated with a given disorder) are located at the top of the list. We then tested whether a given gene set is over-represented at the top of the list by performing an incremental enrichment analysis (**Methods**, see **Supplementary Table 4** for a full GO result). This approach allows us to (1) identify biological pathways associated with a given trait regardless of the power of GWAS, and (2) characterize the biological pathways reflecting the gene set as a whole rather than using arbitrarily defined genes with a specific P-value threshold.

All brain disorders showed enrichment for pathways involved in transcriptional and translational regulation (e.g. transcriptional regulators, RNA splicing, and DNA damage and repair pathways; Table 1). This is in line with previous findings that transcriptional dysregulation may mediate risk for brain disorders^27,28^. Neuronal differentiation and neuronal apoptotic pathways were also enriched in all brain disorders. Neurogenesis was enriched in the majority of disorders except ASD and BD, consistent with an increasing number of studies elucidating the role of neurogenesis, differentiation, and neuronal apoptosis in brain disorders^29–31^. Not surprisingly, neurotransmitter and synaptic pathways were implicated in multiple brain disorders, supporting decades of studies highlighting the importance of synaptic function in psychiatric disorders^32–34^.

**Table 1.** Gene ontologies enriched for brain disorders.

|  | ADHD | ASD | BD | SCZ | MDD | AD | PD | MS | ALS |
| --- | --- | --- | --- | --- | --- | --- | --- | --- | --- |
| Transcriptional regulators | ✓ | ✓ | ✓ | ✓ | ✓ | ✓ | ✓ | ✓ | ✓ |
| DNA damage/repair | ✓ | ✓ | ✓ | ✓ | ✓ | ✓ | ✓ | - | ✓ |
| RNA splicing | ✓ | ✓ | ✓ | ✓ | ✓ | ✓ | ✓ | ✓ | ✓ |
| Neurogenesis | ✓ | - | - | ✓ | ✓ | ✓ | ✓ | ✓ | ✓ |
| Neuronal differentiation | ✓ | ✓ | ✓ | ✓ | ✓ | ✓ | ✓ | ✓ | ✓ |
| Neural apoptosis | ✓ | ✓ | ✓ | ✓ | ✓ | ✓ | ✓ | ✓ | ✓ |
| Glutamatergic | ✓ | ✓ | ✓ | ✓ | ✓ | ✓ | ✓ | ✓ | ✓ |
| GABAergic | - | ✓ | ✓ | ✓ | - | - | - | - | ✓ |
| Synaptic | Pre/Post | Post | Post | Pre/Post | Pre/Post | Post | Post | Pre/Post | Post |
| Neurotransmitter | Monoamine, Dopamine, Acetylcholine, Serotonin | Acetylcholine, Serotonin | Acetylcholine | Acetylcholine, Dopamine | Monoamine | - | Norepinephrine | Dopamine | - |
| Glia/Astrocytes | ✓<br>Differentiation | ✓<br>Development, Gliogenesis | ✓<br>Gliogenesis | - | ✓<br>Development, Migration | ✓<br>Differentiation, Proliferation, Migration | ✓<br>Differentiation, Proliferation | ✓<br>Proliferation, Projection, Migration | ✓<br>Migration |
| Oligodendrocytes | - | ✓ | - | ✓ | ✓ | ✓<br>(Myelination) | ✓<br>(Myelination) | ✓<br>(Myelination) | ✓<br>(Myelination) |
| Brain development | Cortex | Telencephalon | Telencephalon, Hippocampus, Substantia Nigra | Cortex | Telencephalon, Cortex, Substantia Nigra | Cortex, Forebrain | Telencephalon, Forebrain, Limbic system, Midbrain | Cortex, Forebrain, Hypothalamus | Forebrain, Hindbrain, Substantia Nigra |
| Interferon | ✓ | - | ✓ | ✓ | ✓ | ✓ | ✓ | ✓ | ✓ |
| Antigen processing | ✓ | ✓ | ✓ | ✓ | - | ✓ | ✓ | ✓ | ✓ |
| Interleukin | ✓ | ✓ | ✓ | ✓ | ✓ | ✓ | ✓ | ✓ | ✓ |
| Cytokine | - | ✓ | - | ✓ | ✓ | ✓ | ✓ | ✓ | - |
| T cell | ✓ | ✓ | ✓ | ✓ | ✓ | ✓ | ✓ | ✓ | ✓ |
| B cell | - | ✓ | - | ✓ | - | ✓ | - | - | - |
| TLR | ✓ | ✓ | ✓ | ✓ | ✓ | - | ✓ | - | ✓ |
| Aging | ✓ | ✓ | - | - | - | ✓ | - | - | ✓ |
| Ischemia | - | - | - | - | - | - | - | ✓ | ✓ |
| Others | ERBB | BAF, APP, Wnt, Vocalization | APP, Glucocorticoid | Amyloid beta, Wnt, β-catenin, ERBB, BAF | β-catenin, ERBB | Amyloid beta, ERBB, Glucocorticoid | Amyloid beta, Wnt, ERBB, Tau | Wnt, β-catenin, ERBB, Tau | Wnt |

There were interesting distinctions among brain disorders. For example, all brain disorders showed postsynaptic associations, while a selected set of disorders (ADHD, SCZ, MDD, and MS) also exhibited presynaptic associations. Further, while the majority of brain disorders displayed enrichment in glutamatergic signaling, ASD, SCZ, and ALS displayed enrichment in GABAergic signaling. ASD-associated genes were also enriched for acetylcholinergic and serotonergic signaling, reflecting known biology of ASD^35,36^. SCZ and BD-associated genes were also enriched for acetylcholinergic signaling, supporting previous studies that altered cholinergic signaling contributes to SCZ and BD pathogenesis^4,37^. MS-associated genes were enriched for dopaminergic signaling, disruption of which has been associated with immune malfunction in MS^38^. These results collectively highlight synaptic dysfunction in brain disorders, albeit we could detect distinctions among disorders based on neurotransmitters and pre/post synaptic associations.

We observed pronounced immune response for degenerative disorders in contrast to psychiatric disorders. In support of this finding, multiple aspects of glial development were also associated with brain disorders, with stronger enrichment in neurodegenerative disorders (Table 1). This is in line with the heritability enrichment, from which neurodegenerative disorders exhibited enrichment in blood cells (Figure 2A). Immune-related processes and microglia, the brain’s primary immune cells, have been shown to play vital roles in neurodegeneration^21,39^. Moreover, all neurodegenerative disorders showed associations with genes involved in myelination and oligodendrocyte function, suggesting a potential role of oligodendrocytes in neurodegeneration. In line with this, single cell transcriptomic profiles in AD postmortem brains suggested altered molecular profiles in oligodendrocytes^40^. Together with heritability enrichment, this finding of enriched immune response in degenerative, but less so in psychiatric disorders hints a possible explanation for genetic distinctions between psychiatric and degenerative disorders.

Additional interesting findings include amyloid beta enrichment for AD and PD and tau enrichment for MS and PD (Table 1), supporting the amyloid beta and tau pathology in neurodegenerative disorders^41,42^. We also observed Wnt/β-catenin pathway enrichment for a number of brain disorders including ASD, SCZ, MDD, PD, MS and ALS. Wnt/β-catenin signaling is a key pathway for neurogenesis and cortical pattern specification and its dysregulation has been observed in several psychiatric disorders^43–45^. Notably, genes involved in vocalization were associated with ASD, diagnostic criteria of which include impairment in vocalization^46^. We also identified brain regions (e.g. cortex, hippocampus, substantia nigra, and hypothalamus) associated with multiple brain disorders. This is intriguing given that we used cortical, but not subcortical Hi-C data. Psychiatric disorders in general were enriched for cortical genes, while neurodegenerative disorders also showed enrichment for multiple sub-cortical regions, also pointing a potential divergence between psychiatric and neurodegenerative disorders based on brain circuits.

### Cell-type specificity

Identification of the critical cell types for brain disorders is essential for developing proper therapeutic strategies. Indeed, brain disorders often exhibit different cellular signatures and vulnerability. For example, autism postmortem brains exhibit cell-type specific gene expression signatures such as upregulation of glial genes and downregulation of neuronal genes^47^. Common variation in SCZ maps onto specific groups of cells including pyramidal neurons and medium spiny neurons^48^. Microglia are increasingly recognized as a central cell type contributing to the etiology of AD^20^.

To address central cell types mediating risk for brain disorders, we next assessed cell-type specific expression profiles of brain disorder-associated genes (**Methods**). One striking difference between psychiatric and neurodegenerative disorders was that psychiatric disorder-associated genes coalesced in neurons, while neurodegenerative disorder-associated genes were highly expressed in glia (microglia for AD and MS, astrocytes for ALS) except PD (Figure 4A). This finding is in line with the known microglia-specific vulnerability of AD and MS^17–20^. Since psychiatric disorders showed neurodevelopmental origin (Figure 2), we also measured cell-type specific expression profiles of psychiatric disorder-associated genes in the developing cortex (**Methods**). Psychiatric disorder-associated genes converged onto outer radial glia and excitatory neurons (Figure 4B). This selective enrichment in excitatory neurons prevailed across development, as adult neuronal expression profiles for psychiatric disorder-associated genes also indicated excitatory neuronal enrichment (Figure 4B). Collectively, psychiatric disorders coalesced at the level of cellular expression profiles, while degenerative disorders showed distinct expression profiles, implying pleiotropy in psychiatric disorders may stem from shared biology.

**Figure 4.**
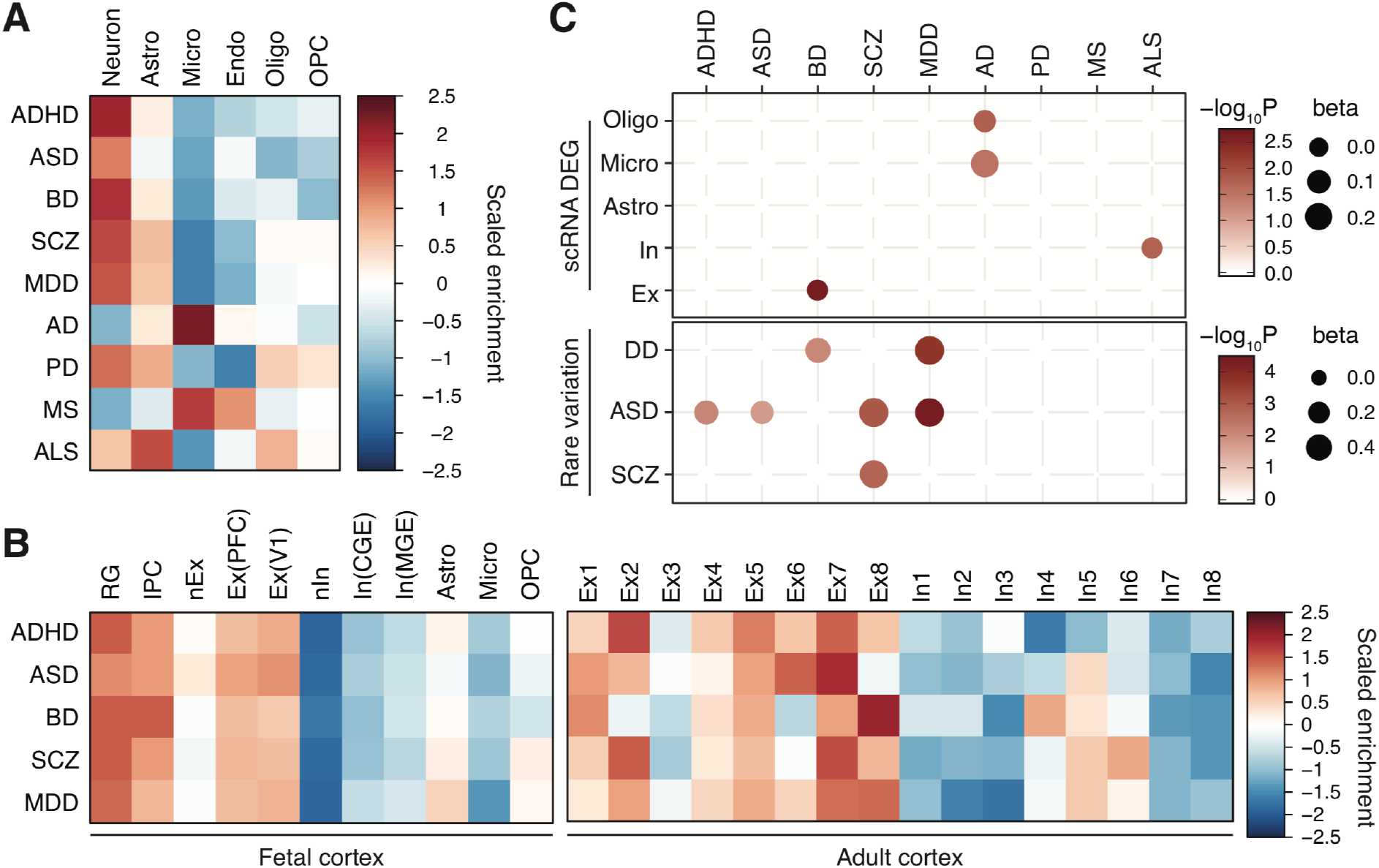
Characteristics of brain disorder-associated genes. **A.** Psychiatric disorder-associated genes are highly expressed in neurons, while neurogenerative disorder-associated genes exhibit glial signatures. Astro, astrocytes; Micro, microglia; Endo, endothelial cells; Oligo, oligodendrocytes; OPC, oligodendrocytes progenitor cells. **B.** Psychiatric disorder-associated genes are highly expressed in radial glia and excitatory neurons. RG, radial glia, vRG; ventricular RG; oRG, outer RG; tRG, truncated RG; IPC, intermediate progenitor cells; Ex, excitatory neurons; In, inhibitory neurons; nEx/nIn, newly born excitatory/inhibitor neurons; PFC, prefrontal cortex; V1, visual cortex; CGE, caudal ganglionic eminence; MGE, medial ganglionic eminence. **C.** Comparison between common variation-associated genes and cell-type specific differentially expressed genes for AD (scRNA DEG, top) and rare variation harboring genes (bottom). Only significant associations (FDR<0.1) were depicted.

### Functional impact of genetic risk factors in transcriptomic signatures

We next hypothesized that brain disorder-associated genes are dysregulated in corresponding disorders. Therefore, we assessed whether brain disorder-associated genes are differentially regulated in postmortem brains with brain disorders. Since different brain disorders have different numbers of significantly associated genes (Figure 1B), we tried to avoid selecting genes based on an FDR threshold. Instead, we used the gene-set analysis in MAGMA which utilizes the whole gene-level association statistics controlling for covariates such as gene size and linkage disequilibrium (LD, **Methods**)^7^.

We first compared our gene association statistics with postmortem brain gene expression profiles from individuals with three psychiatric disorders (ASD, BD, SCZ)^49^. We found a significant overlap between common variation affected genes and differentially expressed genes (DEG) in SCZ (**Supplementary Figure 3A**). SCZ-associated genes were also enriched for ASD DEG. In addition, ADHD-associated genes were enriched for ASD DEGs, recapitulating a shared genetic relationship between these two neurodevelopmental disorders (Figure 3A, **Supplementary Figure 3A**).

Since gene co-expression networks may capture additional disease-associated signatures to DEG, we then compared gene association statistics with gene co-expression modules in three psychiatric disorders (**Supplementary Figure 3B**). We found that an interneuronal module (geneM23) downregulated in ASD were enriched for ADHD-associated genes. Further, SCZ-associated genes were enriched for a synaptic module (geneM7) upregulated in BD and SCZ. MDD-associated genes were enriched for a neurodevelopmental module (geneM16) upregulated in BD and SCZ.

However, the overlap between DEG and gene association statistics was nominal (beta 0-0.04), and ASD- and BD-associated genes were neither differentially regulated in psychiatric disorders nor enriched in any disease-associated co-expression modules (**Supplementary Figure 3**). This can be due to multiple reasons. First, ASD and BD GWAS have relatively limited power compared to SCZ and MDD, hence a more comprehensive picture may arise once we obtain better powered GWAS. Second, transcriptomic signatures do not necessarily reflect early events in the disease process that are directly impacted by common genetic components, but result from complex gene-environment interactions throughout the disease progression. Third, given that brain disorder-associated genes show cell-type specific enrichment (Figure 4A), they may affect gene regulation in a specific cell type(s) that may be missed by the bulk expression datasets. To test the third hypothesis, we compared gene association statistics with cell-type specific molecular signatures in AD pathology^40^. We found that AD-associated genes were significantly enriched for DEGs in microglia and oligodendrocytes, but not in neurons (Figure 4C). While we cannot completely rule out the first and second hypotheses, this result suggests that the cellular context in which risk variants influence gene expression needs to be carefully considered in understanding the molecular complexity of brain disorders.

### Interplay between common and rare variation

Not only common but also rare variation plays a role in brain disorders, highlighting the importance of studying risk variants across the allele frequency spectrum. We have previously shown that genes affected by common SCZ-associated variants are enriched for genes that harbor rare protein disrupting variation in ASD^50^, suggesting a potential interplay between common and rare variation in the genetic architecture of brain disorders. Therefore, we assessed how rare and common variation in brain disorders crosstalk with each other at a gene level using the gene-set analysis in MAGMA (**Methods**).

We found that SCZ-associated genes are enriched for genes that harbor rare *de novo* variation in SCZ and ASD (**Supplementary Figure 4**). This is in line with our previous findings that rare and common variation in SCZ may impact same biological pathways^50^. Genes that are affected by both common (H-MAGMA FDR<0.05) and rare variation in schizophrenia include synaptic genes (*DLG2*, *SYNGAP1*, *SHANK1*) and transcriptional regulators (*SETD1A*, *SMARCC2*). In addition, MDD-associated genes showed enrichment with genes that harbor rare *de novo* variation in ASD and DD. This result suggests that a recently reported genetic correlation between MDD and ASD GWAS (Figure 3A)^51^ may also apply for rare variation (**Supplementary Figure 4**). Another interesting finding was the association between AD-associated genes with genes with rare *de novo* variation in DD, suggesting that genes involved in intellectual capacity may be shared between neurodevelopmental and neurodegenerative disorders.

The gene-set analysis in MAGMA does not take exon length, but gene length, into account, while the number of rare variation is dependent on the exon length. Therefore, we used an alternate approach to control for exon length and the number of SNPs mapped to the genes as covariates (**Methods**). Most of the findings were recapitulated in this framework except an association between AD-associated genes and DD risk genes (Figure 4C). Moreover, ADHD- and ASD-associated genes were also enriched for genes with rare *de novo* variation in ASD, supporting shared genetic basis of neurodevelopmental disorders.

### Biological pathways underlying pleiotropy

Shared genetic etiology across psychiatric disorders (Figure 2) may underlie concerted developmental expression trajectories and cellular expression profiles of psychiatric disorder-associated genes. Cross-disorder GWAS meta-analyzing eight types of psychiatric disorders recently identified more than a hundred GWS loci increasing risk for multiple disorders, indicating widespread pleiotropy among psychiatric disorders^24^. Therefore, we next examined genes shared in multiple psychiatric disorders (n≥3) to identify common molecular mechanisms of psychiatric disorders (see **Methods** for gene selection).

In total, we found 2,772 genes (hereby referred to as pleiotropy genes) that are shared in more than three psychiatric disorders. As expected, these genes strongly overlapped with putative target genes for meta-analysis GWAS of eight psychiatric disorders (OR=7.47, CI=6.11-9.09, P=2.42×10^−68^)^24^, suggesting that they may have shared genetic basis. Pleiotropic genes were involved in gene regulation, neuronal activation, and synaptic and dendritic development (Figure 5A). They showed a distinct peak at mid-gestation, which was consistent with the overall developmental expression patterns of psychiatric disorder associated genes (Figure 2B, 5B). Finally, pleiotropic genes showed strong neuronal enrichment, which was biased toward excitatory neurons compared to inhibitory neurons (Figure 5C). Notably, they were highly expressed in excitatory neuronal subtypes 7-8, corticothalamic projection neurons in cortical layer 6. They also showed relative enrichment in excitatory neuronal subtype 1, cortical projection neurons in cortical layers 2/3.

**Figure 5.**
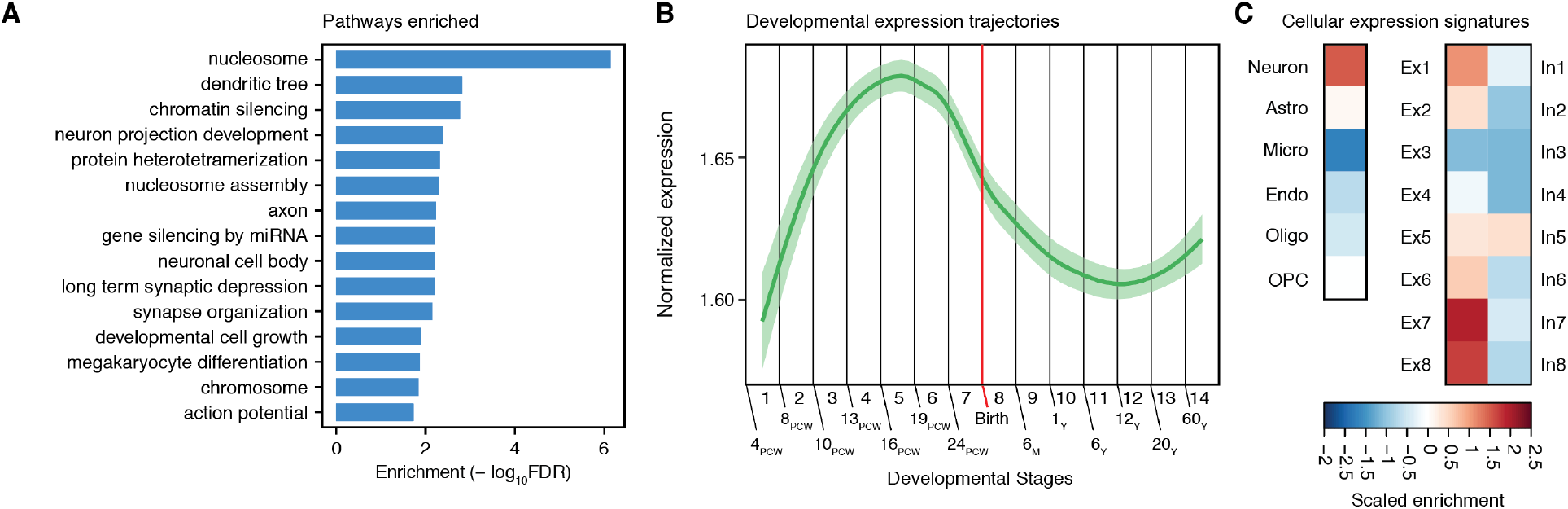
Pleiotropic genes reveal shared molecular mechanisms of psychiatric disorders. **A.** Gene ontology enrichment of pleiotropic genes. **B.** A developmental expression trajectory of pleiotropic genes. LOESS smooth curve with 95% confidence bands. **C.** Cell-type specific expression profiles of pleiotropic genes.

## Discussion

Here we present a refined framework for gene pathway analysis, H-MAGMA, that aggregates SNP-level summary statistics into the gene-level association statistics. Compared with conventional MAGMA, H-MAGMA (1) gains power by linking non-coding SNPs to their target genes based on physical proximity, and (2) adds relevant cellular context to gene mapping by using chromatin interaction data from appropriate tissue and cell types. While the basic concept of mapping SNPs to genes using functional genomic resources is similar to FUMA^14^, H-MAGMA leverages the MAGMA framework to obtain gene-level association statistics in a genome-wide fashion, while FUMA maps a selected set of genomic loci to target genes. Therefore, H-MAGMA can provide an attractive alternative to compare different GWAS to identify shared pathways and identify genes and biological pathways for low powered GWAS.

This framework can be expanded into many different forms. For example, while we decided to use MAGMA among many other tools as it is most widely used, this framework can be applicable to any other tools that convert SNP-level P-values into gene-level association statistics. Moreover, this framework can be readily applicable using different Hi-C datasets to generate gene-level association statistics of non-brain GWAS (e.g. Hi-C datasets from immune cells for rheumatoid arthritis). Although we primarily used Hi-C datasets to link SNPs to target genes, other functional genomics approaches such as eQTLs^13,52^, chromatin accessibility correlations^53^, and machine learning-based enhancer-promoter predictions^54^ can be used to generate SNP-gene pairs.

An application of this framework to nine brain disorder GWAS enabled systematic delineation of pathogenic mechanisms of brain disorders. For example, one important question in psychiatry is whether a critical window exists for treatment of psychiatric disorders. Moreover, it is an ongoing debate whether adult onset disorders such as schizophrenia and depression have a neurodevelopmental origin. By comparing prenatal and postnatal expression trajectories of the brain disorder-associated genes, we found that genes associated with psychiatric disorders show prenatal enrichment, while genes associated with neurodegenerative disorders show postnatal enrichment. In particular, psychiatric disorder-associated genes displayed remarkable developmental convergence onto the mid-gestation, suggesting that shared genetic risk factors for psychiatric disorders may exert their pathogenicity during this period. Mid-gestation is the period during which neurogenesis for upper layer neurons, dendritic arborization and axonogenesis, and initial synaptogenesis takes place^23^. In contrast, neurodegenerative disorder-associated genes were gradually increased upon age, reflecting their increased burden upon aging.

Another layer of convergence among psychiatric disorders was hinted by cellular expression profiles. Psychiatric disorder-associated genes were selectively expressed in excitatory neurons, while neurodegenerative disorder-associated genes show more diverse cellular enrichment profiles. For example, AD- and MS-associated genes showed enhanced expression in microglia, while ALS-associated genes were highly expressed in astrocytes. PD-associated genes were highly expressed in neurons and astrocytes. Similar cell-type specificity was reported by the cell-type specific interactome study, demonstrating the robustness of result using orthogonal approaches^55^.

These results collectively suggest that the shared genetic basis of psychiatric disorders translates into shared neurobiological mechanisms. To further identify shared neurobiological basis among psychiatric disorders, we defined a set of pleiotropy genes that are associated with more than three psychiatric disorders. Pleiotropy genes were enriched for genes involved in neuronal activity and synaptic plasticity, suggesting that inappropriate neuronal activity regulation may act as a key component in the pathogenesis of psychiatric disorders. Pleiotropy genes also displayed mid-gestational and excitatory neuronal enrichment, which summarizes the overall pattern of psychiatric disorder-associated genes. Importantly, this characteristic was also observed for pleiotropic genes identified by meta-analysis of eight psychiatric disorders^24^.

Finally, we found intricate relationships among genes impacted by common and rare variation. For example, common and rare variation in SCZ coalesce to the same set of genes, and common variation in SCZ affect the genes that also harbor rare variation in ASD. Recently observed genetic relationships between ASD and MDD^2^ was detected at the level of rare and common variation. This result highlights the importance of studying risk variation across the allele frequency spectrum to comprehensively understand the complex interplay between common and rare variation in psychiatric disorders.

## Acknowledgement

We thank members of the Won lab, Geschwind lab, and Daniel H. Geschwind for helpful discussions and comments about this paper. This research was supported by a National Institute of Mental Health grant (R00MH113823, H.W.), a NARSAD Young Investigator Award from the Brain and Behavior Research Foundation (H.W.), a SPARK grant from the Simons Foundation Autism Research Initiative (H.W.), training grants from the UNC Neuroscience Center (5T32NS007431, N.Y.A.S., H.F.), and Helen Lyng White Fellowship (W.M.).

## Author Contributions

H.W. designed the H-MAGMA framework. N.Y.A.S., H.F., W.M. conducted the analyses. N.Y.A.S. and H.W. wrote the manuscript.

## Methods

### GWAS

We used the following GWAS summary datasets:

| GWAS | Reference |
| --- | --- |
| Attention deficit/hyperactivity disorder (ADHD) | 56 |
| Autism spectrum disorder (ASD) | 57 |
| Bipolar disorder (BD) | 58 |
| Schizophrenia (SCZ) | 59 |
| Major depressive disorder (MDD) | 60 |
| Alzheimer's disease (AD) | 61 |
| Parkinson's disease (PD) | 62 |
| Multiple sclerosis (MS) | 63 |
| Amyotrophic lateral sclerosis (ALS) | 64 |

### H-MAGMA gene annotation

Exonic and promoter SNPs were directly assigned to their target genes based on their genomic location. The gene model was based on Gencode v26, after converting exon coordinates from hg38 to hg19 using `LiftOver`^65^. Promoters were defined as 2kb upstream of transcription start sites (TSS) of each gene isoform. Intronic and intergenic SNPs were assigned to their cognate genes based on chromatin interactions. Promoter- and exon-based interaction profiles were defined as previously described^13,66^. Briefly, we generated a background Hi-C interaction profile by pooling 9 million imputed SNPs from schizophrenia GWAS summary statistics^67^. Using this background Hi-C interaction profile, we fit the distribution of Hi-C contacts at each distance from each chromosome using the `fitdistrplus` package (https://cran.r-project.org/web/packages/fitdistrplus/index.html). Significance for a given Hi-C contact was calculated as the probability of observing a stronger contact under the fitted Weibull distribution matched by chromosome and distance. Hi-C contacts with FDR<0.01 were selected as significant interactions. Significant Hi-C interacting regions were overlapped with Gencode v.26 exon and promoter coordinates to identify exon- and promoter-based interactions. We generated MAGMA input files using fetal and adult brain Hi-C, in which genes and their assigned SNPs were defined (**Supplementary Data File**). Input files can be also found in the github repository: https://github.com/thewonlab/H-MAGMA.

### Running MAGMA

For conventional MAGMA, we used the MAGMA analysis pipeline as the default setting:

~~~
magma_v1.07b/magma --bfile g1000_eur –pval <GWAS summary statistics> use=rsid,p ncol=N --gene-annot snp_gene.genes.annot --out <output file>
~~~

Here, `g1000_eur` and `snp_gene.genes.annot` denote the reference data file for European ancestry population and the gene location file, respsectively. Both files can be downloaded from: https://ctg.cncr.nl/software/magma.

For H-MAGMA, we used the identical setting except using H-MAGMA inputs guided by Hi-C interaction profiles.

~~~
magma_v1.07b/magma --bfile g1000_eur --pval <GWAS summary statistics> use=rsid,p ncol=N --gene-annot <H-MAGMA input annotation file> --out <output file>
~~~

Detailed instructions can be found in the github repository: https://github.com/thewonlab.

### Comparison between H-MAGMA and conventional MAGMA

Since H-MAGMA uses Ensemble gene IDs, while conventional MAGMA uses Entrez gene ID, we converted Entrez gene IDs to Ensemble gene IDs using `biomaRt`, then merged MAGMA outputs from H-MAGMA and conventional MAGMA using the same gene identifiers. Only genes that were detected in both measures were selected, and their Z-scores from H-MAGMA and conventional MAGMA were compared by Pearson correlation. Pearson correlation coefficients were plotted to visualize the extent of similarities between two frameworks.

### Partitioned heritability enrichment

To measure heritability enrichment of eight brain disorder GWAS in active genomic regions in each cell/tissue-type, we used stratified LDSC with the baseline-LD model (S-LDSC)^16^ and chromHMM-defined chromatin states^15^. Since chromatin profiling hasn’t been performed in all cell/tissue-types (e.g. DNase hypersensitivity was missing for fetal brains, while H3K27ac ChIP-seq was not performed in the adult DLPFC), we instead used genomic regions that are active in each cell/tissue type using chromatin states defined by chromHMM^68^. We defined active genomic regions by the regions that are marked as Active transcription start sites (state 1), Flanking active TSS (state 2), Genic enhancers (state 6), and Enhancers (state 7) in the core 15-state model (https://egg2.wustl.edu/roadmap/web_portal/chr_state_learning.html). The SNP annotation file can be downloaded from the github repository: https://github.com/thewonlab/H-MAGMA. Heritability enrichment values in different cell/tissue types resulting from the `S-LDSC` were then scaled to allow tissue-level comparison of enrichment values.

### Gene selection

For assessing (1) developmental expression profiles, (2) cell-type specific expression profiles, and (3) gene ontology enrichment of disorder-associated genes, we used following strategies to select genes. We restricted our analysis only on protein-coding genes, because (1) the majority of genes detected in the spatiotemporal transcriptomic atlas^22^, single cell expression datasets^69–71^, and gene ontologies were protein-coding genes, and (2) non-coding genes have much lower expression values compared with protein-coding genes, which can dilute the signals. We also excluded genes within the MHC region due to the complexity of LD, which can override the overall pattern. Finally, we removed genes within chromosome X (chrX), as only a subset of GWAS had association statistics available in chrX.

### Developmental and Cellular Expression Profiles

Analyzing developmental and cell-type specific expression levels required selection of significantly associated genes for each disorder. We calculated adjusted P-values based on the Benjamini and Hochberg (BH) procedure using `p.adjust` function in `R.` We then selected genes with two FDR thresholds (FDR<0.01 for GWAS with >20 GWS hits, SCZ, BD, MDD, AD; FDR<0.1 for GWAS with < 20 GWS hits, ADHD, ASD, PD, MS, ALS) for as significantly associated brain disorder genes.

We used spatiotemporal transcriptomic atlas from Kang et al., 2011^22^. We used cortical expression profiles across multiple developmental stages. Log-transformed expression values were centered to the mean expression level per sample using a `scale(center=T, scale=F)` function in `R`. Genes associated with brain disorders were selected for each brain sample and their average centered expression values were calculated for each brain sample. Since H-MAGMA results were available from fetal and adult brain Hi-C, we used genes that are significantly associated in either fetal or adult brain using a `union` function in `R`. Prenatal versus postnatal expression values were compared using `lm` function function in `R` (e.g. for a given disorder, `lm(Expression values ~ stages)`).

Developmental stages were defined as below:

| Developmental Stage | Age |
| --- | --- |
| 1 | 4 PCW ≤ Age < 8 PCW |
| 2 | 8 PCW ≤ Age < 10 PCW |
| 3 | 10 PCW ≤ Age < 13 PCW |
| 4 | 13 PCW ≤ Age < 16 PCW |
| 5 | 16 PCW ≤ Age < 19 PCW |
| 6 | 19 PCW ≤ Age < 24 PCW |
| 7 | 24 PCW ≤ Age < Birth |
| 8 | Birth ≤ Age < 6 M |
| 9 | 6M ≤ Age < 1 Y |
| 10 | 1 Y ≤ Age < 6 Y |
| 11 | 6 Y ≤ Age < 12 Y |
| 12 | 12 Y ≤ Age < 20 Y |
| 13 | 20 Y ≤ Age < 60 Y |
| 14 | Age > 60 Y |

We also used single cell transcriptomic data from the adult brain^69,71^ and fetal brains^70^ to identify cell-type specific expression profiles of brain disorder-associated genes. We also processed log-transformed expression values per cell using a `scale(center=T, scale=F)` function in R. Average centered expression values of genes associated with brain disorders were calculated for each cell type. We used H-MAGMA results derived from fetal brain Hi-C to assess cell-type specific expression values in the fetal brain^70^. H-MAGMA results derived from adult brain Hi-C were used to assess cell-type specific expression values in the adult brain^69,71^.

### Gene ontology

We used an `R` package `gProfileR` (https://biit.cs.ut.ee/gprofiler/gost) for running gene ontology analysis, as it allows using a ranked gene list, which resembles Gene Set Enrichment Analysis (GSEA, http://software.broadinstitute.org/gsea/index.jsp). This does not require P-value threshold to select significantly associated genes, which allows comparing gene ontologies for differently powered GWAS in a non-bias fashion. We used Hi-C derived H-MAGMA results from both fetal and adult brains for psychiatric disorders and adult brain Hi-C derived H-MAGMA results for neurodegenerative disorders. After ranking genes based on Z-scores generated by H-MAGMA, we ran gene ontology analysis using this command line:

~~~
gprofiler(<Ranked gene list>, organism="hsapiens", ordered_query=T, significant=T, max_p_value=0.05, min_set_size=15, max_set_size=600, min_isect_size=5, correction_method="fdr", hier_filtering="moderate", custom_bg=background gene set, include_graph=T, src_filter="GO")
~~~

### Gene-set analysis

Genes that harbor *de novo* protein disrupting variation in developmental disorders (DD) were obtained from the Deciphering Developmental Disorders Study (93 DD risk genes with genome-wide significance)^72^. We also obtained 99 ASD risk genes (rare variation burden, FDR<0.1) from the Autism Sequencing Consortium (ASC) study^73^. Schizophrenia risk genes with elevated burden of rare variation were obtained from Singh et al., 2016^74^ (110 genes with FDR<0.3). Differentially expressed genes (DEG) in postmortem brains with psychiatric disorders were obtained from Gandal et al., 2018^49^ (FDR<0.05). Cell-type specific DEG in Alzheimer’s disease brains was obtained from Mathys et al., 2019^40^. Using these gene sets, we used the gene-set analysis in MAGMA with this command line:

~~~
magma --gene-results <MAGMA outputfile.genes.raw> --set-annot <Rare variation annotation file> --out <Output filename>
~~~

Given that the protein disrupting variation in brain disorders was detected by exome-sequencing and is dependent on the exon, but not gene length, we also used an alternate approach to compare MAGMA outputs (common variation) with the gene lists that harbor protein disrupting variation (rare variation). Here we regressed out the exome length (controlling for rare variation) and the number of SNPs mapped to each gene using MAGMA (controlling for common variation).

~~~
glm(<MAGMA Z-scores> ~ <Rare variation annotation file> + <Exome length>
+ <The number of SNPs mapped to genes>)
~~~

### Rank-rank hypergeometric overlap (RRHO)

We assessed genetic relationship between two disorders (r_g_) by using genetic correlation analysis of LDSC^1^. To provide similar metrics based on gene-level association statistics, we compared ranks between two datasets (e.g. MAGMA outcomes from two disorders) using `R` package `RRHO` (https://www.bioconductor.org/packages/release/bioc/html/RRHO.html) with the following command line:

~~~
RRHO(<Ranked gene list 1>, <Ranked gene list 2>, outputdir=<output directory>, alternative="enrichment", BY=TRUE, log10.ind=TRUE)
~~~

To compare gene-level overlaps (RRHO output) with genetic correlations (calculated by LDSC), P-values from RRHO was converted into Z-scores using the following command line:

~~~
Zscore = qnorm(10^(-Pvalues), lower.tail=FALSE)
~~~

We then compared resulting RRHO Z-scores with r_g_ values from the genetic correlation analysis using Pearson’s correlation. This correlation coefficient provides a metric to compare genetic relationship between two disorders measured at the genome-wide SNP level (r_g_) vs. gene-level (RRHO Z).

### Identification of pleiotropic genes

RRHO outputs two gene sets consisting of most up and downregulated genes, with most upregulated genes referring to a list of genes that are associated with both conditions, and most downregulated genes referring to a list of genes that are not associated with both conditions. Therefore, we employed most upregulated genes as a gene list that is shared between two disorders, hence marking pleiotropic genes. We then generated pleiotropic genes shared in at least three disorders by intersecting RRHO most upregulated genes between the following disorder pairs (ADHD vs. ASD/BD/SCZ/MDD; ASD vs. BD/SCZ/MDD; BD vs. SCZ/MDD; and SCZ vs. MDD). We next generated a merged gene set by a union function in R and obtained uniquely identified genes. The code is provided in the github repository: https://github.com/thewonlab/H-MAGMA. In the end, we obtained 2,772 genes that are shared in more than three disorders, and defined them as pleiotropic genes. We next performed the gene ontology, developmental expression, and cell-type expression analyses on the pleiotropic genes as previously described.

### Data availability

All GWAS summary statistics used in this study are publicly available. We deposited (1) H-MAGMA input files derived from the fetal brain and adult brain Hi-C data, and (2) H-MAGMA output files for nine brain disorders in the github repository https://github.com/thewonlab/H-MAGMA.

### Code availability

Codes used in this study are provided in the github repository: https://github.com/thewonlab/H-MAGMA

