## Supplementary tables for "Connecting gene regulatory relationships to neurobiological mechanisms of brain disorders"

**Supplementary Table 1.** H-MAGMA outputs for brain disorders based on fetal and adult brain Hi-C data.

**Supplementary Table 2.** Comparison between the genes identified by H-MAGMA and conventional MAGMA.

**Supplementary Table 3.** Statistical comparison between prenatal and postnatal expression values for each brain disorder-associated genes.

**Supplementary Table 4.** Gene ontologies of brain disorders based on fetal and adult brain H-MAGMA.

**Supplementary Table 5.** A list of pleiotropic genes.

### Supplementary Figures

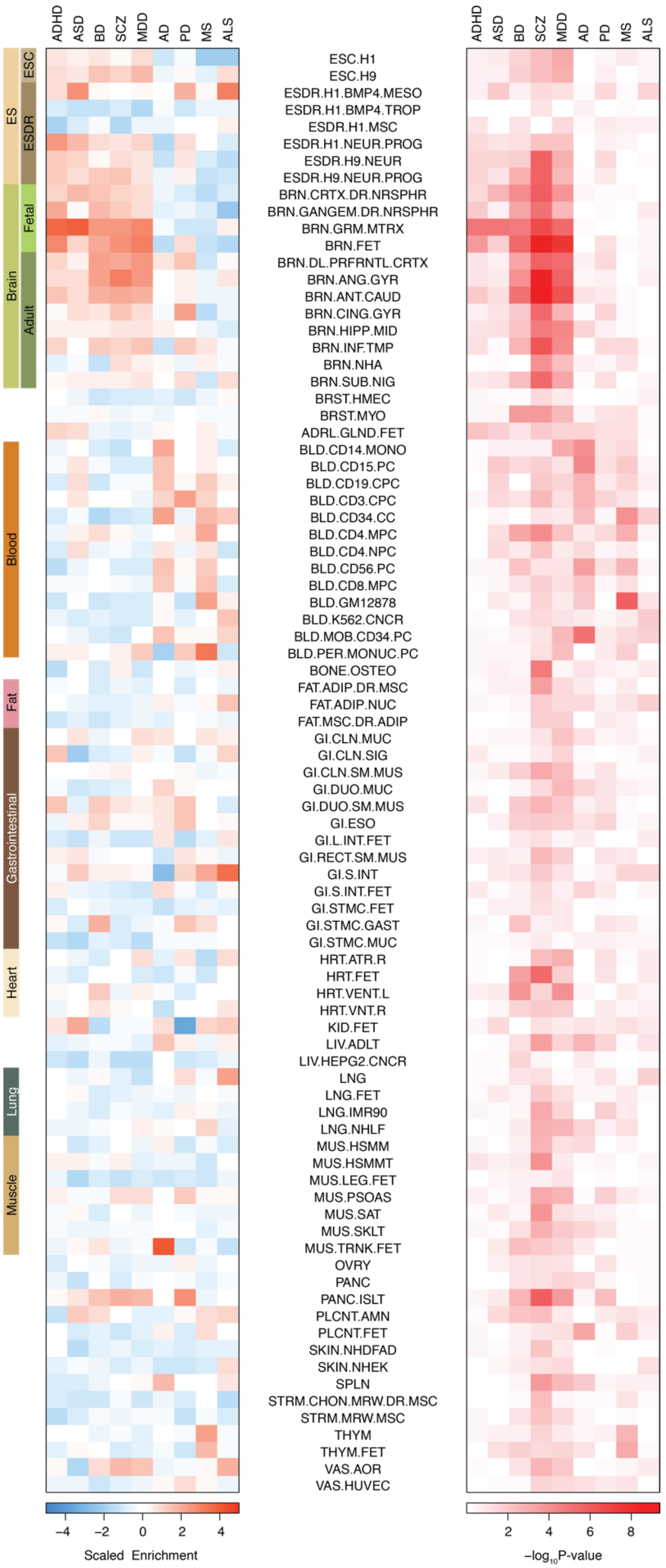


**Supplementary Figure 1. Heritability enrichment of brain disorders in active regulatory elements of multiple tissue/cell types.** Scaled enrichment values are plotted in the top, and significance of heritability enrichment (P-values) are plotted in the bottom. ESC, embryonic stem cells. ESDR, embryonic stem cell derived cell lines.

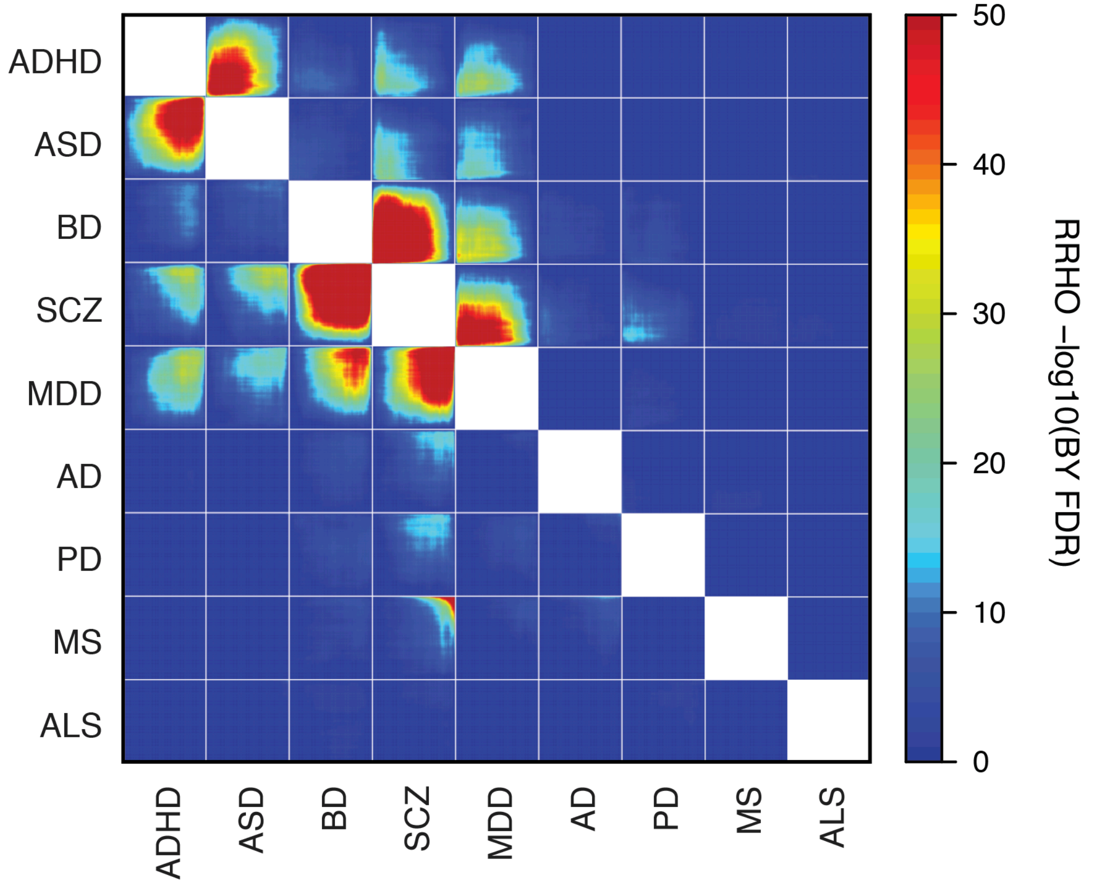


**Supplementary Figure 2. Genetic relationships between nine brain disorders based on gene-level overlaps. Top right,** Rank-rank hypergeometric overlaps (RRHO) of H-MAGMA outputs from the fetal brain. **Bottom left,** RRHO of H-MAGMA outputs from the adult brain.

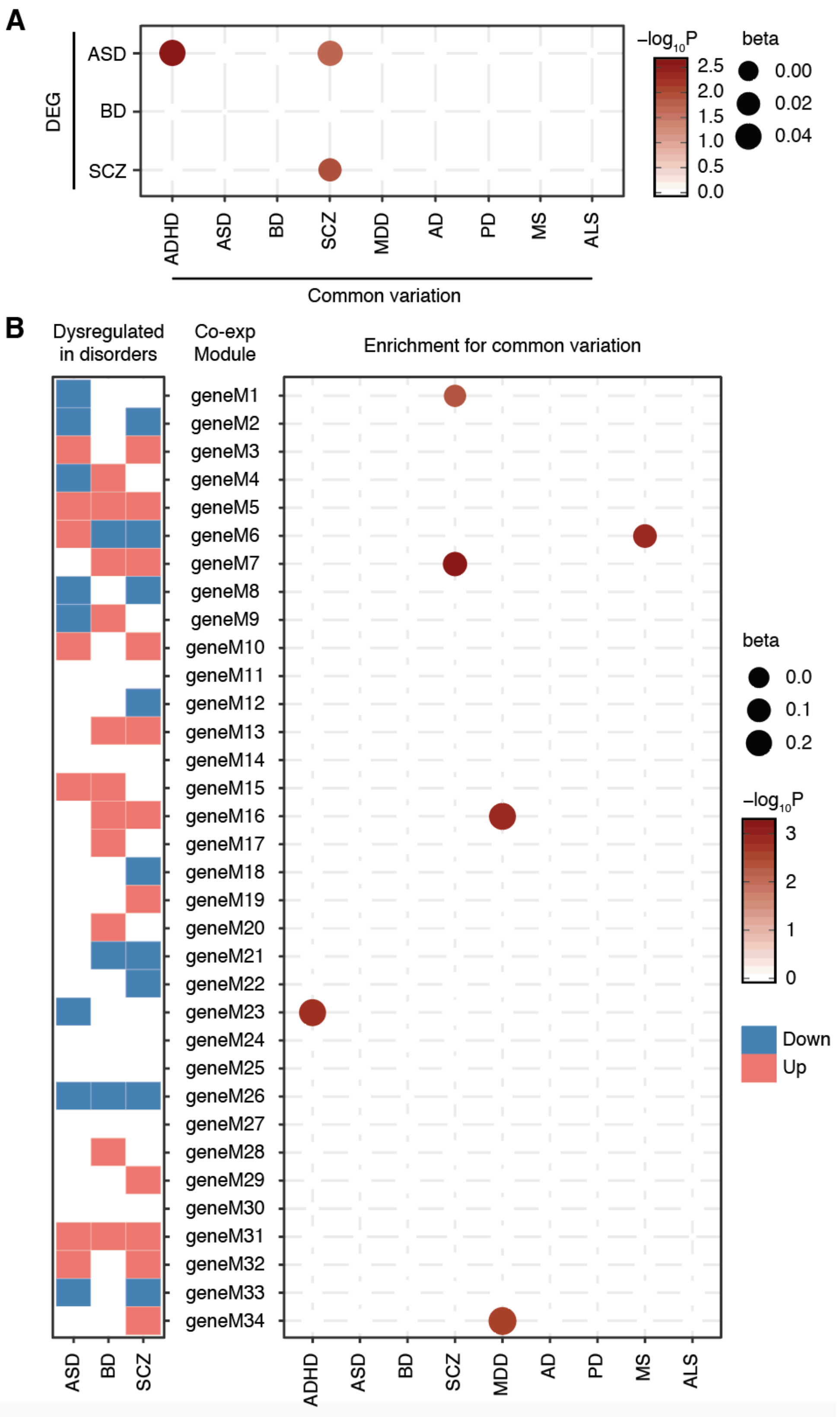


**Supplementary Figure 3. Common variation-associated genes are dysregulated in postmortem brains of individuals with psychiatric disorders. (A).** Overlap between common-variation associated genes and genes differentially expressed (DEG) in postmortem brains with psychiatric disorders. **(B).** Overlap between common-variation associated genes and co-expression (co-exp) modules differentially regulated in psychiatric disorders. Down, modules are downregulated in disorders; Up, modules are upregulated in disorders.

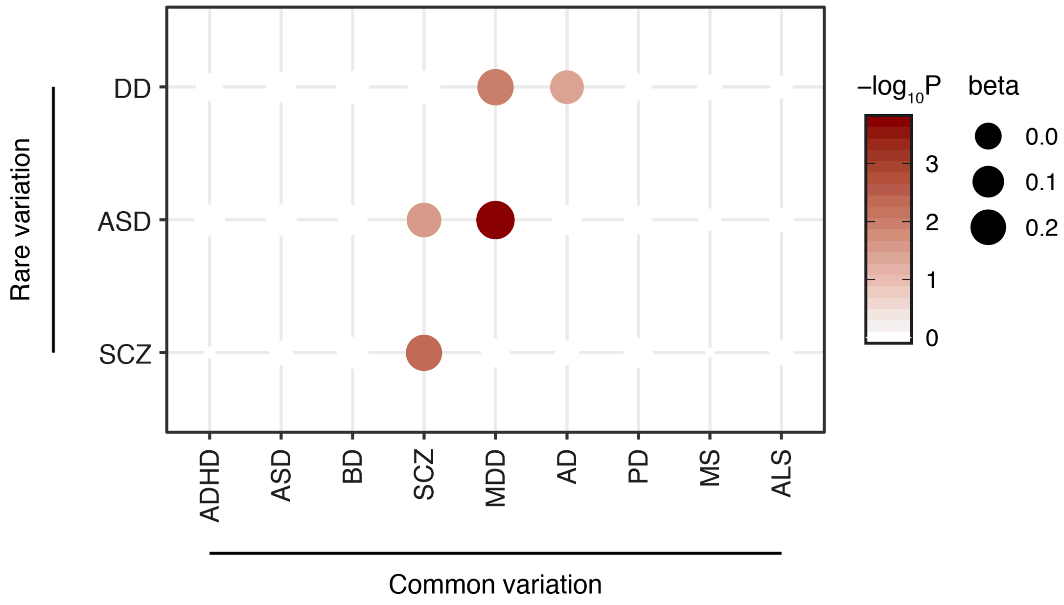


**Supplementary Figure 4. Overlap between common variation-associated genes and rare variation harboring genes.** Enrichment scores based on the MAGMA-based gene-set analysis, which corrects for gene length, minor allele frequency, and gene density. Compare with the result in Figure 4C, which depicts the same enrichment result after correction for exon length and SNP numbers. DD, developmental disorders.
